# *Cutibacterium acnes* clonal complexes display various growth rates in blood-culture bottles used for diagnosing orthopedic device-related infections

**DOI:** 10.1101/2021.01.11.426311

**Authors:** Faten El Sayed, Petra Camernik, Anne-Laure Roux, Lea Papst, Thomas Bauer, Lionelle Nkam, Valérie Sivadon-Tardy, Latifa Noussair, Jean-Louis Herrmann, Jean-Louis Gaillard, Samo Jeverica, Martin Rottman

**Author notes:** <u>Corresponding author</u>: Faten El Sayed, @.

## Abstract

Blood-culture bottles (BCBs) are widely used to improve the diagnosis of orthopedic device-related infections. Data is scarce on the growth of *Cutibacterium acnes* and its genotypes in BCBs under real-life clinical conditions.

We studied 39 cases of revision arthroplasty for which at least one intraoperative sample yielded a pure *C. acnes* culture from anaerobic BCBs (BD Bactec Lytic/10 Anaerobic/F [Lytic Ana]) and/or solid media. Genotyping of *C. acnes* isolates from the 39 cases allowed: i) the identification of 49 non-redundant isolates belonging to four clonal complexes (CCs): CC18, CC28, CC36, and CC53 and ii) the determination of infectant and contaminant strains. Under real-life clinical conditions, Lytic Ana alone was more often positive for contaminants than infectant strains (18/36 [50%] *versus* 2/13 [15.4%]; p = 0.047). The time to detection (TTD) values in Lytic Ana were shorter for CC53 than other CCs (mean [SD] TTD: 77 [15] *versus* 165 [71] hours; p = 0.02). CC53 was confirmed to grow faster than other CCs by studying an enlarged panel of 70 genotyped *C. acnes* strains inoculated *in vitro* into Lytic Ana vials (mean [SD] TTD: 73 [13] *versus* 122 [50] hours; p < 0.001).

The use of Lytic Ana BCBs in orthopedics increases the recovery rate of *C. acnes* but leads to the isolation of proportionally more contaminants than true infectant strains. TTD values are much shorter for CC53 strains, irrespective of their being infectant or contaminant. TTD does not solely reflect the bacterial load of samples but also clonal complex-related traits.

## INTRODUCTION

*Cutibacterium acnes* is an anaerobic aero-tolerant microorganism that is involved in orthopedic device-related infections (ODRIs) and is a frequent cause of shoulder prosthetic joint infection. The diagnosis of *C. acnes* ODRI is, however, challenging in clinical practice. *C. acnes* ODRIs are often associated with few clinical manifestations and normal or subnormal levels of inflammatory markers. It is a common inhabitant of the human skin and sebaceous glands and may be a contaminant. Finally, it is a fastidious organism that is difficult to isolate from clinical samples.

Recent advances have been made in the microbiological diagnosis of ODRIs by inoculating relevant samples (e.g. synovial fluid or homogenates of periprosthetic tissue) into aerobic and anaerobic blood culture bottles (BCBs), which are then incubated and monitored in an automated device (1–5). However, data is scarce on the relevance of this approach for the diagnosis of ODRIs caused by *C. acnes*. Minassian et al. found that 96% of anaerobic cultures, including those of *C. acnes*, were detected within five days, and 99% within 10 days using the BD Bactec system. However, incubating anaerobic BCBs beyond seven days was shown to yield only contaminants, all *C. acnes* (2). Similarly, we previously showed that “infectant” *C. acnes* were all detected within seven days of incubation when anaerobic BCBs were inoculated with bead-milled tissue samples (unpublished data).

The previous studies, including ours, did not focus specifically on *C. acnes* and included results from only a small number of *C. acnes* strains. Recently, Rentenaar *et al.* studied the detection of a panel of 26 clinical *C. acnes* isolates in various BCBs. However, the experiments were performed by *in vitro* inoculation of BCBs and were mainly aimed at comparing the performance of BCB vials available in the Becton Dickinson system (6). Moreover, there was no information about the genotypes of the isolates.

“Orthopedic” strains inoculated *in vitro* into anaerobic BCBs were recently found to be associated with a broad range of TTD values (7). This prompted us to study whether such variability may be related to the genotype and to evaluate its clinical impact.

## PATIENTS AND METHODS

### Background information

The surgery department of the Ambroise Paré hospital is a French reference center for the management of bone and joint infections. All patients undergoing surgery for ODRIs or suspicion of an ODRI have at least three intraoperative samples taken for microbiological culture using both BCBs and solid media.

### Bacteriological methods

Intraoperative tissue samples were bead-milled in sterile water (8) and the homogenates inoculated: i) onto solid media and incubated for five days (Columbia sheep blood agar under aerobic and anaerobic conditions and chocolate agar under 5% CO_2_), ii) into aerobic BCBs (BD Bactec Peds+, Becton Dickinson Diagnostics, Sparks, MD) and incubated for seven days, and iii) into anaerobic BCBs (BD Bactec Lytic/10 Anaerobic/F; “Lytic Ana”) and incubated for 14 days. Aerobic and anaerobic BCBs were monitored in the Bactec FX instrument (Becton Dickinson Diagnostics) and subcultured only if the instrument gave a positive result. Bacterial isolates were identified by mass spectrometry using a Microflex LT instrument and the current CE-marked IVD Biotyper software (Bruker Daltonique, Wissenbourg, France). Subsequent cryopreservation was performed on colonies randomly selected from a pure subculture plate.

### *C. acnes* genotyping

Cryopreserved *C. acnes* isolates were genotyped using a previously described multi-locus sequence typing (MLST) scheme (9). Purified PCR products were sequenced using the BigDye® Terminator v1.1 kit on an Applied Biosystems 3500xl Dx sequencer. The sequence type (ST) was determined using a publicly available MLST database (http://pacnes.mlst.net). The clonal complex (CC) was determined by eBURST analysis (eBURST version 2 - http://eburst.mlst.net/).

### Definitions

A *C. acnes* strain was considered to be infectant if it was cultured from at least two distinct samples (same ST and same antibiotic susceptibility pattern) belonging to the same case or a contaminant if it was cultured from only one sample. A *C. acnes* case was considered to be an infection if at least one infectant *C. acnes* strain was recovered. Otherwise, it was considered to be a contamination.

If two or more samples were positive for the same ST in a given patient, only one sample, selected at random, was considered for further analyses, under both clinical and experimental conditions.

### Study under real-life clinical conditions

We included 39 cases for which: i) at least one intraoperative sample yielded a pure *C. acnes* culture from Lytic Ana and/or anaerobic solid media, ii) *C. acnes* isolates were genotyped, and iii) no other microbial pathogen was recovered. Recovery rates on each culture media and the time to detection (TTD) value of *C. acnes* isolates in Lytic Ana were retrieved from the laboratory information software.

### Study under experimental *(in vitro)* conditions

In addition to the isolates recovered from the 39 cases, we added 20 additional cases selected according to the same clinical criteria but for which the growth data from the initial culture could not be retrieved from the laboratory information software to enlarge our collection.

A 0.5-McFarland suspension of *C. acnes,* was prepared from cryopreserved isolates subcultured for 72 h on Columbia 5% sheep blood agar. Serial dilutions in sterile saline were performed to obtain ~10^2^ colony-forming units (CFUs)/mL. Two “Lytic Ana” vials without blood or additives were inoculated with 1 mL (~10^2^ CFUs). The inoculum density and viability were controlled by enumeration of CFUs on Columbia 5% sheep blood agar under anaerobic conditions. Vials were monitored in a Bactec FX for 14 days or until positivity. Positive vials were subcultured on aerobic and anaerobic Columbia sheep blood agar to verify culture purity.

### Data and Statistical analyses

Univariate analysis was performed using Fisher’s exact test for categorical variables and the Wilcoxon-Mann Whitney test for continuous variables. A p-value < 0.05 was considered significant for all statistical analyses.

## RESULTS

### See flowchart in supplementary file for a summary of cases and strains included in clinical and experimental conditions

#### Cases and strains included in the “clinical conditions” study

The patients for the 39 cases studied were mostly male (25/39, 64.1%). The cases included 13 of *C. acnes* infection and 26 of *C. acnes* contamination; three cases harbored both infectant and contaminant *C. acnes* strains and were classified as “infections”. The patients for cases of infection were younger than those for cases of contamination, were less likely to have had surgery for revision arthroplasty, and more likely to have had spinal surgery (Table 1).

**Table 1.**
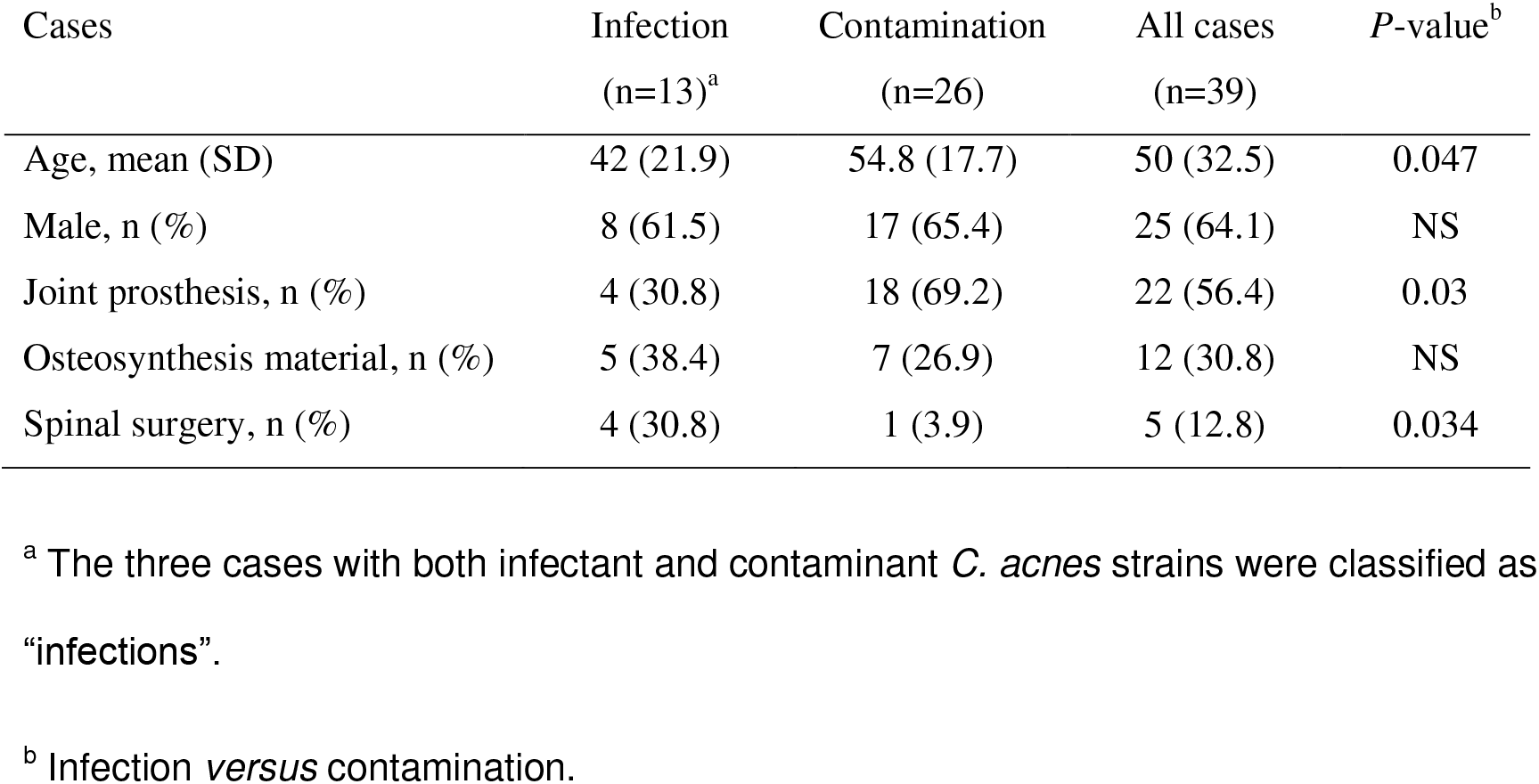
Characteristics of cases.

| Cases | Infection<br>(n=13)<sup>a</sup> | Contamination<br>(n=26) | All cases<br>(n=39) | *P*-value<sup>b</sup> |
| --- | --- | --- | --- | --- |
| Age, mean (SD) | 42 (21.9) | 54.8 (17.7) | 50 (32.5) | 0.047 |
| Male, n (%) | 8 (61.5) | 17 (65.4) | 25 (64.1) | NS |
| Joint prosthesis, n (%) | 4 (30.8) | 18 (69.2) | 22 (56.4) | 0.03 |
| Osteosynthesis material, n (%) | 5 (38.4) | 7 (26.9) | 12 (30.8) | NS |
| Spinal surgery, n (%) | 4 (30.8) | 1 (3.9) | 5 (12.8) | 0.034 |

<sup>a</sup> The three cases with both infectant and contaminant *C. acnes* strains were classified as “infections”.
<sup>b</sup> Infection *versus* contamination.

In total, 49 non-redundant *C. acnes* isolates (“*C. acnes* strains”) were recovered from the 39 cases (one ST, 31 cases; 2 STs, 6 cases; 3 STs, 2 cases) and included 13 infectant and 36 contaminant strains. The 49 strains were distributed among four CCs, with the most frequent being CC36 (38.8%), followed by CC18 and CC53 (22.4% each), and CC28 (16.3%); CC18 (38.5%) was the most frequent CC among infectant strains and CC36 (44.4%) among contaminants (Table 2). The overall distribution of CCs showed, however, no significant difference between infectant strains and contaminants (p = 0.36, NS).

**Table 2.** Genotypes of strains recovered from the cases.

| Strains | % (nb) of strains |  |  |  |
| --- | --- | --- | --- | --- |
|  | CC18 | CC28 | CC36 | CC53 |
| All strains (n=49) | 22.4 (11) | 16.3 (8) | 38.8 (19) | 22.4 (11) |
| Infectant (n=13) | 38.5 (5) | 15.4 (2) | 23.1 (3) | 23.1 (3) |
| Contaminants (n=36) | 16.7 (6) | 16.7 (6) | 44.4 (16) | 22.2 (8) |

#### Recovery of strains and clonal complexes from Lytic Ana*versus* solid media under clinical conditions

No *C. acnes* was ever recovered from PedsPlus vials. The included strains were only slightly more often recovered from Lytic Ana than solid media (63.3% [31/49] *versus* 59.2% [29/49]). Overall, strains of *C. acnes* were more often recovered from Lytic Ana alone (40.8%) or solid media alone (36.7%) than from both (22.5%) (Table 3). The recovery rates differed, however, among infectant strains and contaminants: Lytic Ana and solid media together were more often positive for infectant strains (53.8% *versus* 11.1% with contaminants, p = 0.004), whereas Lytic Ana was more often positive alone for contaminants (50% *versus* 15.4% with contaminants, p = 0.047).

**Table 3.**
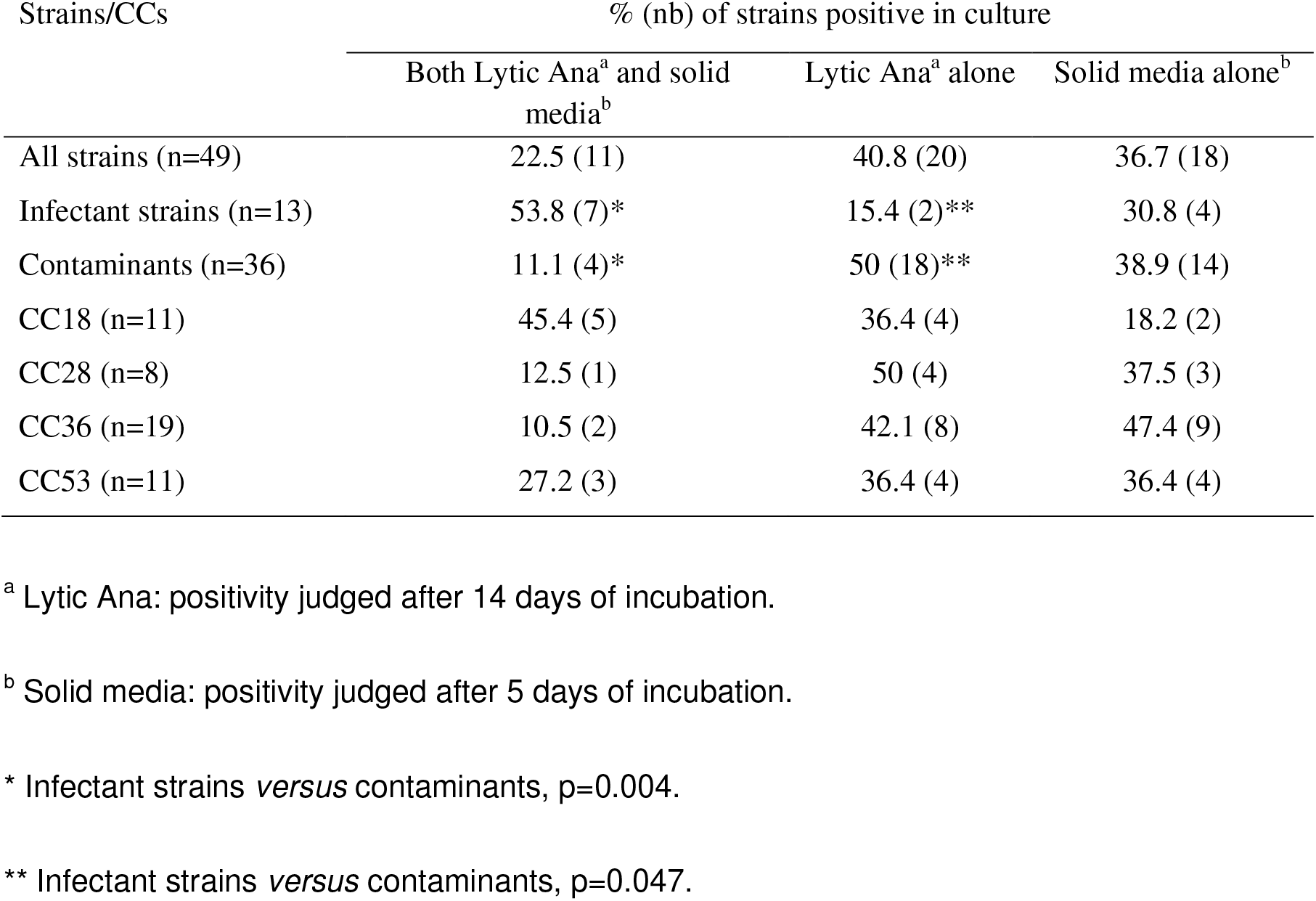
Recovery of strains and CCs from Lytic Ana and solid media.

| Strains/CCs | % (nb) of strains positive in culture |  |  |
| --- | --- | --- | --- |
|  | Both Lytic Ana<sup>a</sup> and solid media<sup>b</sup> | Lytic Ana<sup>a</sup> alone | Solid media alone<sup>b</sup> |
| All strains (n=49) | 22.5 (11) | 40.8 (20) | 36.7 (18) |
| Infectant strains (n=13) | 53.8 (7)* | 15.4 (2)** | 30.8 (4) |
| Contaminants (n=36) | 11.1 (4)* | 50 (18)** | 38.9 (14) |
| CC18 (n=11) | 45.4 (5) | 36.4 (4) | 18.2 (2) |
| CC28 (n=8) | 12.5 (1) | 50 (4) | 37.5 (3) |
| CC36 (n=19) | 10.5 (2) | 42.1 (8) | 47.4 (9) |
| CC53 (n=11) | 27.2 (3) | 36.4 (4) | 36.4 (4) |

<sup>a</sup> Lytic Ana: positivity judged after 14 days of incubation.
<sup>b</sup> Solid media: positivity judged after 5 days of incubation.
\* Infectant strains *versus* contaminants, p=0.004.
\*\* Infectant strains *versus* contaminants, p=0.047.

The recovery rates of CC18, CC28, C36, and CC53 were 81.8%, 62.5%, 52.6%, and 63.6%, respectively, for Lytic Ana, and 63.6%, 50%, 57.9%, and 63.6%, respectively, for solid media. There was no significant difference in the distribution of CCs among strains recovered from the various media (Table 3).

#### Lytic Ana TTD values of infectant strains and contaminants under clinical and experimental conditions

TTD values were available for 24 of 31 *C. acnes* strains recovered from BCBs. The overall mean (SD) TTD values of *C. acnes* strains (n = 24) in Lytic Ana was 150 (73.2) hours under real-life clinical conditions. The mean values were significantly lower for infectant strains (n = 6) than contaminants (n = 18) (98.3 [53.4] *versus* 167.8 [71.7] hours, p = 0.02) and, at seven days post-inoculation, Lytic Ana was more often positive for infectant strains than contaminants, although the difference did not reach significance (83.3% [5/6] *versus* 38.9% [7/18], p = 0.15) (Fig. 1A). In contrast, the mean TTD values measured under controlled experimental conditions, corresponding to 70 isolates (47/49 + 23), were very similar between infectant strains (n = 23) and contaminants (n = 47) (111 [37.2] versus 115 [54.9] hours, p = 0.67) and the proportion of strains positive at seven days post-inoculation was approximately 80% for both groups (Fig. 1B).

**Figure 1.**
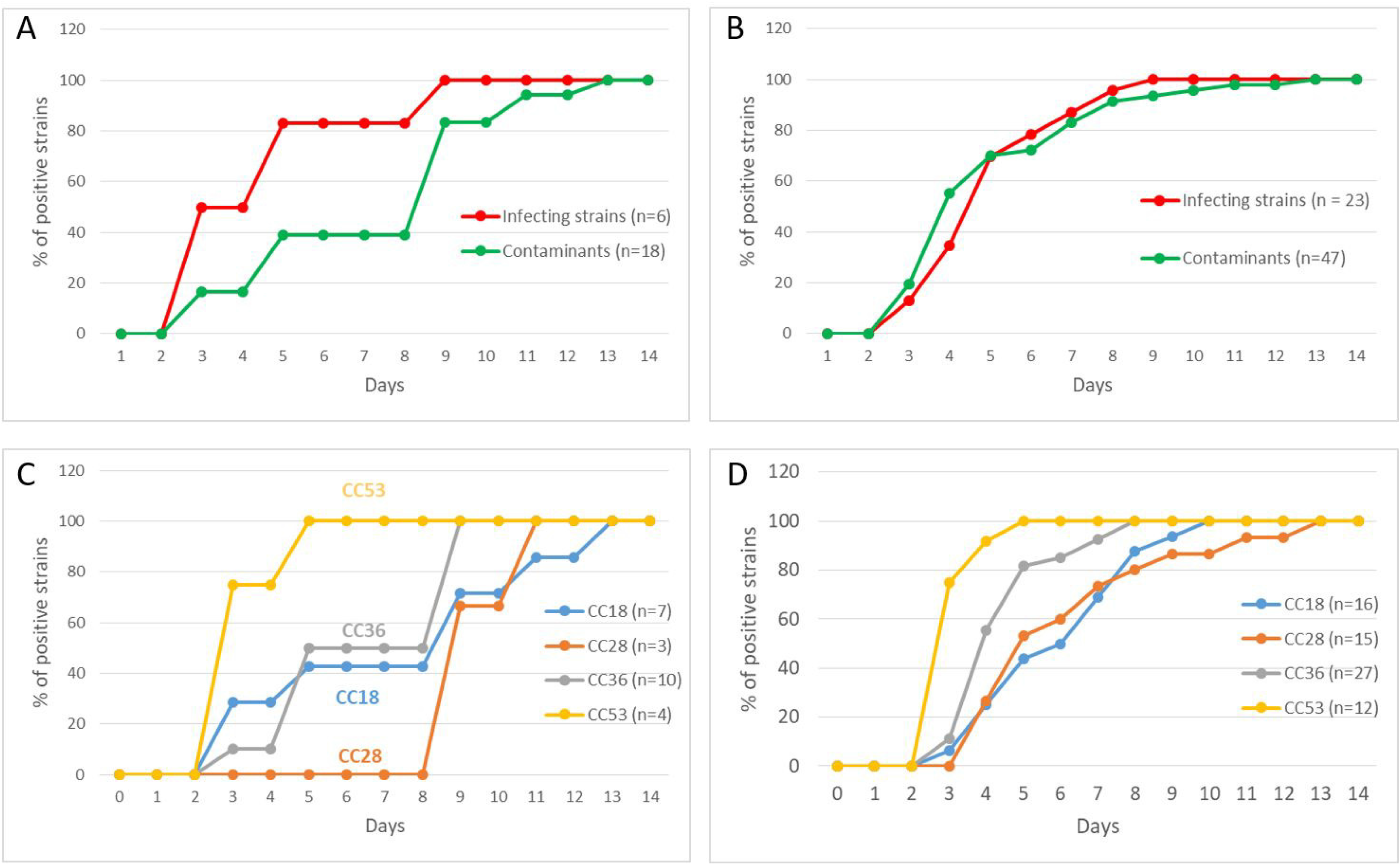
Time to detection of *C. acnes* strains with Lytic Ana under clinical (A, C) and experimental (*in vitro*) conditions (B, D). The cumulative percentage of positive strains for each day for14 days is shown. A, B. Infectant strains *versus* contaminants. C, D. Clonal complexes CC18, CC28, CC36, and CC53.

#### Lytic Ana TTD values of the four main clonal complexes under clinical and experimental conditions

The mean TTD values of the various *C. acnes* CCs measured under clinical conditions ranged from 77.5 (CC53) to 220 hours (CC28) (Table 4). The CC53 strains had significantly lower mean TTD values than all other strains (77.5 [15] *versus* 165 [71] hours, p = 0.02) (Table 4) and were all recovered by seven days post-inoculation (Table 4 and Fig. 1C). The CC36, CC18, and CC28 strains had recovery rates of 50%, 43%, and 0%, respectively, at seven days post-inoculation (Table 4 and Fig. 1C). However, there were only three CC28 strains and all were contaminants.

**Table 4.** TTD values of CCs measured under clinical and experimental conditions.

|  | Clonal complexes |  |  |  | *P*-value<sup>a</sup> |
| --- | --- | --- | --- | --- | --- |
|  | CC18 | CC28 | CC36 | CC53 |  |
| *Clinical conditions* | n=7 | n=3 | n=10 | n=4 |  |
| Mean (SD) TTD values* | 167 [94.5] | 220 [34.6] | 147 [56.6] | 77.5 [15] | 0.02 |
| % (nb) of strains positive at 7 days | 43 (3) | 0 (0) | 50 (5) | 100 (4) |  |
| *Experimental conditions* | n=16 | n=15 | n=27 | n=12 |  |
| Mean (SD) TTD values | 138 [45.8] | 144 [65.2] | 101 [32.9] | 73 [12.8] | <0.001 |
| % (nb) of strains positive at 7 days | 69 (11) | 73 (11) | 93 (25) | 100 (12) |  |

<sup>a</sup> CC53 *versus* all other CCs.
\*TTD values were available for 24 over 31 *C. acnes* strains recovered in blood culture bottles.

The CC53 strains were confirmed to grow faster in Lytic Ana than strains from other CCs under controlled experimental conditions (mean [SD] TTD values: 73 [12.8] *versus* 122 [50] hours, p < 0.001) (Table 4). By seven days post-inoculation, 100% of the CC53 strains had been recovered *versus* 93% of the CC36 strains and approximately 70% of the CC18 and CC28 strains (Table 4 and Fig. 1D).

## DISCUSSION

This is the first study to examine the relationship between *C. acnes* genotype and the time to detection in blood culture vials seeded with intraoperative samples from orthopedic surgery. The blood culture vial used was the Lytic Ana vial, proven for its effectiveness in this setting (6, 7). Nearly 40 cases of infection and contamination with *C. acnes* from our center were included, making it one of the largest clinical series published to date (10–12). Moreover, the MLST typing of all isolates allowed the inclusion of a single ST type per case, limiting redundancy and allowing the rigorous classification of infectant and contaminant cases (13). Finally, this study in a real-life clinical setting was complemented with an *in vitro* study to assay a significant number of isolates from each CC and standardize the inoculum to avoid bacterial load bias.

Our data show that all CCs of *C. acnes* do not grow at the same rate in the Lytic Ana vial. Indeed, CC53 isolates were detected twice as fast as the other CCs, both in real-life clinical settings and *in vitro* in laboratory settings. One hundred percent of CC53 isolates were detected from day 5 after incubation under both clinical and experimental conditions *versus* 50% or less for the other CCs. Moreover, our *in vitro* data suggest that CC36 isolates have an intermediate growth rate between the faster CC53 and slower CC18 and CC28, but these results must be confirmed on a larger dataset. Finally, the abnormally prolonged time to detection of CC28 isolates in the clinical setting was not reproducible *in vitro*, which may be due to the contaminant status of the strains analyzed (see below) or epigenetic imprinting.

We cannot offer a clear explanation for the behavior of CC53 in Lytic Ana vials relative to the other CCs. Nonetheless, CC53 belongs to phylotype II, whereas CC18, CC28, and CC36 belong to phylotype I (9). CC53 isolates may therefore carry metabolic or physiological characteristics that are distinct from the other CC’s that better suit them to growth in Lytic Ana medium, as this trait is not observed in BacT/SN bioMérieux medium (data not shown). The most significant formulation specificity of the Lytic Ana medium is the presence of saponin, a natural detergent composed of an amphipathic glycoside with a lipophilic polycyclic derivative. Saponin may be a direct source of fatty acids that CC53 could specifically use, by analogy with the use of Tween by some mycobacteria or corynebacteria (14–16).

Our study provides several other important observations. First, our data confirm that infective strains of *C. acnes* are detected significantly sooner than contaminants. This difference was entirely abolished when the vials were seeded with a standardized *in vitro*-grown inoculum. The slower growth of contaminant isolates in clinical settings may therefore be associated with a lower bacterial burden. An alternative explanation is the presence of an epigenetic imprint. Our results also show that a growth time between five and eight days discriminates between infectant and contaminant isolates. However, although 60% of contaminant isolates failed to grow within this timeframe, as much as 20% of infective isolates showed delayed growth beyond eight days. This result confirms those of previous studies advocating the extended culture of bone and joint samples beyond eight days when using broth enrichment (e.g., Schaedler broth, brain heart Infusion broth, or thioglycolate broth) (10, 11, 17).

Another important observation is that blood culture vial enrichment is not sufficient for the optimal detection of *C. acnes* in bone and joint infection and that combining it with anaerobic solid media is required. Indeed, as many as 35% of isolates only grew on solid media, whereas 40% of isolates only grew in Ana Lytic medium. Moreover, the recovery rate of contaminants on blood culture media was 50% *vs* 15.4% for infectant strains (p = 0.047), whereas anaerobic solid medium did not significantly favor contaminants (recovery rates of 38.9% *versus* 30.8% for infectant strains). We show that a sample that is simultaneously positive on solid and blood culture media is predictive of infectiveness, with a 53% combined detection rate for infective isolates *vs* 11.1% for contaminants (p = 0.004). Similar results have been reported with thioglycolate, brain heart infusion, and Schaedler broth instead of blood culture vials (10, 11, 17).

Our study had several limitations. In spite of a large number of included isolates, the number of isolates within each CC was limited, which could fail to unveil subtle differences between their characteristics. As mentioned above, CC36 isolates can display an intermediate growth rate in Lytic Ana relative to isolates belonging to CC53 or CCs 18 and 28. Moreover, our data suggest that CC18 isolates grow poorly on agar media, with a recovery rate of 18.2% *vs* 36.4% to 47.4% for the other CCs. We have not tested all STs within each CC and cannot exclude that certain STs would behave differently within a CC. However, this seems unlikely for CC53, which represent a specific phylum within the *C. acnes* population and for which we could evaluate nine different STs.

Our data address solely the Lytic Ana vial and other studies are necessary to determine whether they can be extrapolated to other blood culture media routinely used for the enrichment of bone and joint samples. We have previously reported evidence that each blood culture medium formulation has specific growth characteristics for *C. acnes* (7). The prolonged incubation of liquid media for the enhanced recovery of *C. acnes* has been widely recommended, regardless of its diagnostic value. However, we did not extend the incubation of the CO2-chocolate agar or blood agar anaerobic plates. We do not believe that this negatively affects the detection of pathogens in the context of bead-milled ODRI samples on the basis of two observations: i) we do not routinely observe pinpoint colonies after five days of incubation that would prompt us to perform extended incubations and ii) we did not observe any benefit from incubation past this timepoint when we explored the benefit of prolonged broth incubation or poor sensitivity of agar plate culture that would lead us to question this process.

These results were obtained at a single center and must be confirmed with studies including a larger number of isolates from different regions of the world and multiple centers.

In conclusion, this study is the first to show the impact of the genetic background of *C. acnes* on its growth rate in blood-culture media and further justifies the relevance of molecular typing of *C. acnes*.

## REFERENCES

1. Hughes HC, Newnham R, Athanasou N, Atkins BL, Bejon P, Bowler ICJW. 2011. Microbiological diagnosis of prosthetic joint infections: A prospective evaluation of four bacterial culture media in the routine laboratory. Clin Microbiol Infect.

2. Minassian AM, Newnham R, Kalimeris E, Bejon P, Atkins BL, Bowler ICJW. 2014. Use of an automated blood culture system (BD BACTEC™) for diagnosis of prosthetic joint infections: Easy and fast. BMC Infect Dis.

3. Peel TN, Dylla BL, Hughes JG, Lynch DT, Greenwood-Quaintance KE, Cheng AC, Mandrekar JN, Patel R. 2016. Improved diagnosis of prosthetic joint infection by culturing periprosthetic tissue specimens in blood culture bottles. MBio.

4. Portillo ME, Salvadó M, Trampuz A, Siverio A, Alier A, Sorli L, Martínez S, Pérez-Prieto D, Horcajada JP, Puig-Verdie L. 2015. Improved diagnosis of orthopedic implant-associated infection by inoculation of sonication fluid into blood culture bottles. J Clin Microbiol.

5. Shen H, Tang J, Wang Q, Jiang Y, Zhang X. 2015. Sonication of explanted prosthesis combined with incubation in BD Bactec bottles for pathogen-based diagnosis of prosthetic joint infection. J Clin Microbiol.

6. Rentenaar RJ, Kusen SM, Riemens-Van Zetten GMA, Van Mourika MSM. 2018. Detection of clinical cutibacterium acnes isolates in different becton dickinson blood culture vials. J Clin Microbiol.

7. Jeverica S, El Sayed F, Čamernik P, Kocjančič B, Sluga B, Rottman M, Papst L. 2020. Growth detection of Cutibacterium acnes from orthopaedic implant-associated infections in anaerobic bottles from BACTEC and BacT/ALERT blood culture systems and comparison with conventional culture media. Anaerobe.

8. Roux AL, Sivadon-Tardy V, Bauer T, Lortat-Jacob A, Herrmann JL, Gaillard JL, Rottman M. 2011. Diagnosis of prosthetic joint infection by beadmill processing of a periprosthetic specimen. Clin Microbiol Infect.

9. Lomholt HB, Kilian M. 2010. Population genetic analysis of Propionibacterium acnes identifies a subpopulation and epidemic clones associated with acne. PLoS One 5.

10. Butler-Wu SM, Burns EM, Pottinger PS, Magaret AS, Rakeman JL, Matsen FA, Cookson BT. 2011. Optimization of periprosthetic culture for diagnosis of Propionibacterium acnes prosthetic joint infection. J Clin Microbiol.

11. Bossard DA, Ledergerber B, Zingg PO, Gerber C, Zinkernagel AS, Zbinden R, Achermann Y. 2016. Optimal length of cultivation time for isolation of Propionibacterium acnes in suspected bone and joint infections is more than 7 days. J Clin Microbiol.

12. Frangiamore SJ, Saleh A, Grosso MJ, Alolabi B, Bauer TW, Iannotti JP, Ricchetti ET. 2015. Early versus late culture growth of Propionibacterium acnes in revision shoulder arthroplasty. J Bone Jt Surg - Am Vol.

13. El Sayed F, Roux AL, Sapriel G, Salomon E, Bauer T, Gaillard JL, Rottman M. 2019. Molecular Typing of Multiple Isolates Is Essential to Diagnose Cutibacterium acnes Orthopedic Device-related Infection. Clin Infect Dis.

14. Chevalier J, Pommier MT, Cremieux A, Michel G. 1988. Influence of Tween 80 on the mycolic acid composition of three cutaneous corynebacteria. J Gen Microbiol.

15. Cox FR, Slack CE, Cox ME, Pruden EL, Martin JR. 1978. Rapid tween 80 hydrolysis test for mycobacteria. J Clin Microbiol.

16. Pietersen RD, du Preez I, Loots DT, van Reenen M, Beukes D, Leisching G, Baker B. 2020. Tween 80 induces a carbon flux rerouting in Mycobacterium tuberculosis. J Microbiol Methods.

17. Schäfer P, Fink B, Sandow D, Margull A, Berger I, Frommelt L. 2008. Prolonged Bacterial Culture to Identify Late Periprosthetic Joint Infection: A Promising Strategy. Clin Infect Dis.

